# *MDSvis*, a Shiny application for interactive visualisation of Multi Dimensional Scaling projections

**DOI:** 10.1101/2025.06.23.660991

**Authors:** Philippe Hauchamps, Andrea Vicini, Laurent Gatto

**Affiliations:** Computational Biology and Bioinformatics, de Duve Institute, UCLouvain, Belgium

**Author notes:** Computational Biology and Bioinformatics, de Duve Institute, UCLouvain, Av. Hippocrate 75, 1200 Brussels, Belgium. **Data availability:** The *Krieg_Anti_PD1* cytometry dataset is available from the *HDCytoData* Bioconductor R package. The *fsap* dataset is available from the *MDSvis* GitHub repository: github.com/UCLouvain-CBIO/MDSvis. **Software availability:** The R package is available, under a GPL-3 license, from the *MDSvis* GitHub repository: github.com/UCLouvain-CBIO/MDSvis. The on-line documentation is available, under a CC-BY-4.0 license, from https://uclouvain-cbio.github.io/MDSvis. **Funding:** P.H. is funded by GlaxoSmithKline Biologicals S.A., under a cooperative research and development agreement between GlaxoSmithKline Biologicals S.A. and de Duve Institute (UCLouvain). **Competing interests:** P.H. is a student at the de Duve Institute (UCLouvain) and participates in a post graduate studentship program at GSK. A.V. and L.G. report no competing interest.

## Abstract

**Motivation:** Visual exploratory data analysis is key for making sense of high dimensional data. In particular, Multi Dimensional Scaling (MDS), a low dimensional representation of data points, is frequently used in early stages of data analysis pipelines, for Quality Control (QC), batch effect detection and hypothesis generation. However, interpreting representations in more than two dimensions necessitates the mental integration of multiple two-dimensional (2D) plots - a process that greatly benefits from an interactive Graphical User Interface (GUI), where such plots can be easily and iteratively customised without the need for programming.

**Results:** Here, we present *MDSvis*, a Shiny application for user-interactive customised visualisation of MDS projections. *MDSvis* is based on the Bioconductor package *CytoMDS* for calculating MDS coordinates. We describe the functional workflow of the application, and illustrate two different use cases, which are detailed in the package vignettes. The first use case pertains to cytometry data analysis, which was the original motivation for building the application. The second use case demonstrates how to extend the use of *MDSvis* to MDS visualisation of any sample distance matrix.

**Availability and implementation:** *MDSvis* is implemented in the R programming language and is available in a GitHub repository: *github*.*com/UCLouvain-CBIO/MDSvis*, under a GPL-3 license.

## Introduction

Visual exploratory data analysis is key for interpreting high dimensional data. In particular, low dimensional representations of data points, using e.g. Principal Component Analysis (PCA) [1], or Multi Dimensional Scaling (MDS) [2], are cornerstones of numerous data visualisation and analysis pipelines. Indeed, these multivariate visualisation methods can support early stages of the data analysis process such as Quality Control (QC), batch effect detection and hypothesis generation. In the field of computational flow cytometry [3], we have recently developed *CytoMDS*, a Bioconductor R package for low dimensional representation of single cell cytometry datasets [4]. *CytoMDS* performs fast calculation of Earth Mover’s Distance (EMD) [5] between sample distributions, and generates a metric MDS [6] projection of the different samples, facilitating the detection of outliers and batch effects.

Nevertheless, even in a low dimensional setting, the proper interpretation of representations in more than two dimensions necessitates the cognitive integration of multiple two-dimensional plots - a process that greatly benefits from an interactive Graphical User Interface (GUI), where such plots can be easily and iteratively customised without the need for programming. Hence, providing an interactive visualisation application on top of *CytoMDS*, for exploratory analysis of cytometry datasets, was the original motivation for the current work. However, since MDS has much wider use than cytometry data analysis, we designed the application such that it can also be useful for the MDS representation of any type of sample distances. Here, we present *MDSvis*, a Shiny application [7] enabling users to interactively explore and customise MDS projection plots.

## Software implementation

### Analytical workflow

An overview of the *MDSvis* analytical workflow is shown in Figure 1(A). It can be split into three consecutive steps. The first two steps are performed in an R script, while the last step is done interactively in the *MDSvis* Shiny application. The first step consists in obtaining a sample distance matrix, which can be done in two ways. In the first use case, corresponding to cytometry data analysis, a *CytoMDS* function is called to perform sample distance calculation, using pre-processed cytometry marker expression matrices as input. These matrices typically come from dedicated pre-processing pipelines where cytometry raw data files - in Flow Cytometry Standard (FCS) format - are read, scale transformed and, if applicable, compensated or unmixed. In the second use case, the user uploads a previously generated distance matrix directly. In a second step, in both use cases, *CytoMDS* is called to generate MDS projection results from the input distance matrix. Optionally, the user can also instruct *CytoMDS* to calculate sample statistics, which will be used in bi-plots, for plot interpretation [4]. These results are then saved as R Data Serialize (RDS) objects on disk. In the final step, the RDS objects are uploaded by the user into the *MDSvis* Shiny application, for interactive visualisation. The intermediate storage of the RDS objects allows for asynchronous execution of the visualisation step, hence it can be repeated without re-running the first two steps.

**Figure 1.**
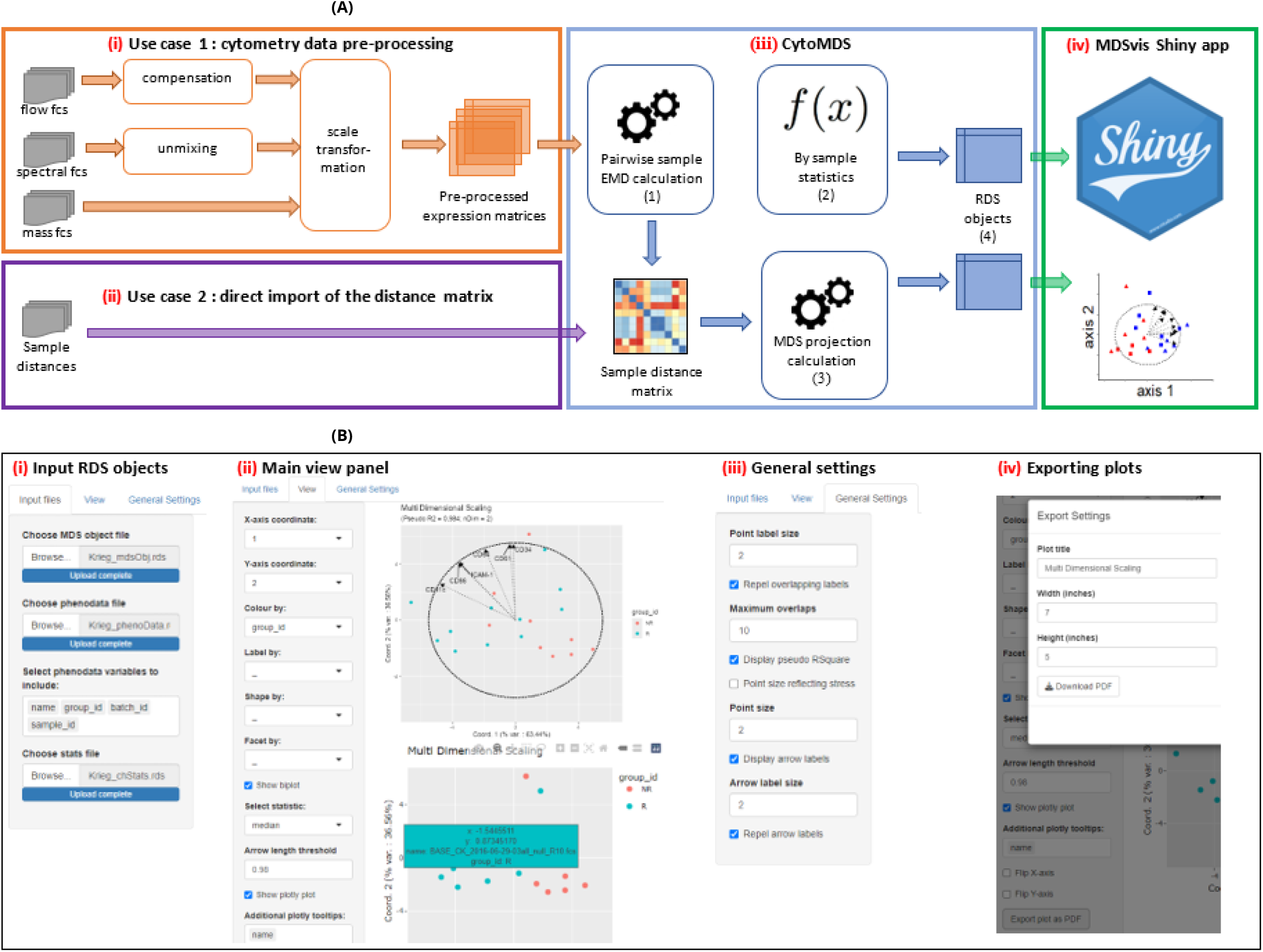
*MDSvis* overview. **(A)** Analytical workflow (figure adapted from [4]). (i) In use case 1, dedicated pre-processing is performed on cytometry data, and the pre-processed expression matrices are then used as input to *CytoMDS* sample distance calculation. (ii) In use case 2, the pairwise sample distance matrix is directly imported. (iii) *CytoMDS* calculation part: in use case 1, pairwise sample EMD distances are calculated from pre-processed cytometry expression matrices (1), and sample statistics can be automatically calculated (2). In both use cases, a MDS projection is calculated from the distance matrix (3). Calculation results are saved as RDS objects (4). (iv) RDS objects are input into the *MDSvis* Shiny application for interactive visualisation. **(B)** *MDSvis* layout and features. (i) Input tab, allowing RDS objects loading. (ii) Main view tab, displaying the MDS projection. Main plot attributes can be customised using sample data characteristics, and an additional *plotly* view can be displayed for zooming into data points. (iii) General plot setting tab. (iv) Export of a generated plot as a file in Portable Document Format (PDF).

#### Application features

Figure 1(B) shows the different tabs of the Shiny application. In the initial ‘Input files’ tab (i), the user is prompted to import the RDS files containing the MDS projection results, sample statistics, and possibly additional sample characteristics - here called *phenoData* - that will be used to customise the plots. Then, the user can switch to the main ‘View’ tab (ii), where the MDS plot is displayed. The user can choose the x and y axes - among the available MDS dimensions. Besides, faceting variable, colour, shape and label of points can be customised by selecting corresponding sample data characteristics from drop-down lists. To facilitate user interpretation, a bi-plot can be overlaid onto the main plot, by selecting the sample statistics to be used - among the ones initially uploaded in the input tab. Finally, an additional interactive *plotly* plot [8] can be activated, which allows to zoom into parts of the plot and display additional data point information. In the ‘General Settings’ tab (iii), secondary plot layout settings, such as size of points, arrows and labels, can be modified by the user. All the generated plots can be exported as files in Portable Document Format (PDF), with customised title and plot dimensions (iv).

## Results

### Cytometry use case

The main *MDSvis* use case is cytometry data analysis - notably QC and batch effect identification. In that case, the *CytoMDS* package is used to calculate sample EMD distances, based on cytometry marker expression matrices. An example of such a use case is described in the on-line package documentation (https://uclouvain-cbio.github.io/MDSvis). In the main vignette, the *Krieg_Anti_PD1* cytometry dataset [9] is used to demonstrate the different steps represented in Figure 1 (use case 1). The dataset consists of 20 baseline mass cytometry samples - prior to treatment - of peripheral blood from melanoma skin cancer patients subsequently treated with anti-PD-1 immunotherapy. The patient samples were split across 2 groups - non-responders and responders - and in 2 acquisition batches. The goal of the study was to identify biomarkers of responsiveness to immunotherapy at baseline. In Figure 1(B), a screenshot of the main *MDSvis* view panel (ii) shows one of several possible visualisations for this dataset - see also Figure 2(A) for a more visible version of this plot. Here, the user selected the first two axes for display and the dots, corresponding to each sample, are coloured according to the two patient groups. The user also chose to overlay a biplot, showing directions of marker medians correlated with the two axes, as well as a unit circle to facilitate quantitative visual assessment of correlations. Finally, a *plotly* interactive plot was activated - Figure 1(B) - for zooming in and displaying - by cursor - some additional sample specific information, like the sample full name, which would be too wide to display as labels on the main plot.

**Figure 2.**
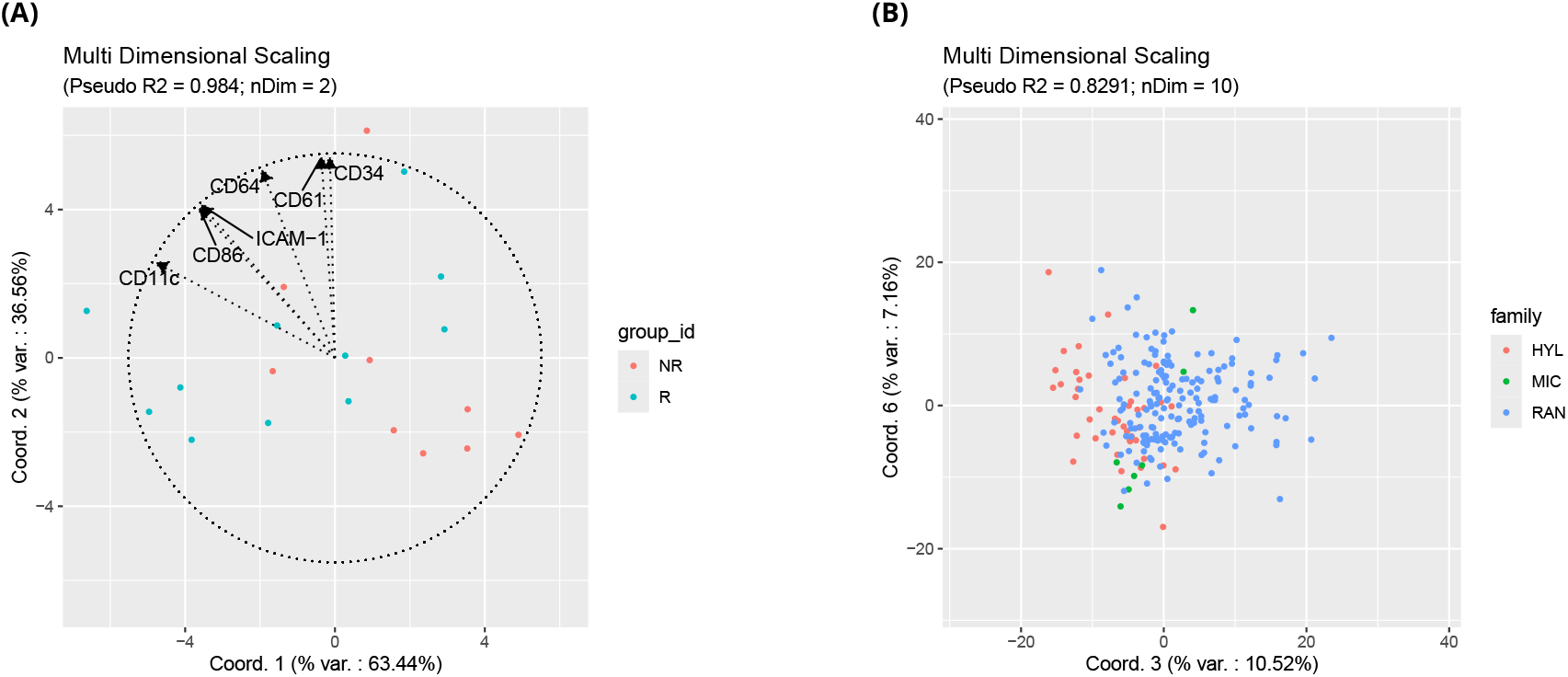
One customised plot of the *Krieg* dataset (A), and one of the *fsap* dataset (B) - exports from *MDSvis*.

### Extension to other MDS applications

The second use case consists in visualising a MDS projection, starting directly from any type of previously obtained sample distance matrix. It is illustrated in a second *MDSvis* package vignette, using the “frog skin active peptide” (fsap) dataset. This dataset consists of 226 samples, where each sample represents an amino acid sequence coding for a skin-secreted host defense peptide from a different species of amphibian (specifically, frogs). An ad-hoc distance matrix between these amino acid sequences was obtained by performing sequence alignment and computing an appropriate dissimilarity indicator (more detailed description is provided in the vignette). Based on this distance matrix, a MDS projection can be calculated by using *CytoMDS* and visualised by using *MDSvis*. One example of plot is shown in Figure 2(B). On that plot, axes 3 and 6 of the projection are represented, and dots are coloured by biological group (family). On that particular view, peptides belonging to the 3 different anuran families overlap while having different plot location distributions.

## Discussion

### Limitation and potential future extensions

The current main limitation of *MDSvis* is the need for the user to initiate the MDS projection calculation programmatically before importing the results into the Shiny application. A future extension could enable users to begin with a precomputed distance matrix and perform the MDS calculation directly within the application. This enhancement would also offer greater flexibility by allowing users to customise aspects of the MDS projection - for example, by selecting a subset of samples to be included in the projection, or, in the context of cytometry, specifying which markers should be used to compute sample distances. Another useful extension, for QC of cytometry samples, would allow the user to dig into a specific sample through dedicated sample views, for instance using *CytoPipelineGUI* [10].

### Conclusion

In this work, we presented *MDSvis*, a Shiny application for interactive customised visualisation of MDS projections, aiming at facilitating data exploration. We described the analytic workflow, as well as the application features, which we illustrated by describing two use cases taken from the package vignettes. We believe that this software will facilitate access to visual exploratory data analysis by lowering the entry barrier, particularly for users with limited programming expertise.

## Declarations

## Data Availability

The *Krieg_Anti_PD1* cytometry dataset is available from the *HDCytoData* Bioconductor R package. The *fsap* dataset is available from the *MDSvis* GitHub repository: github.com/UCLouvain-CBIO/MDSvis.

## Software availability

The R package is available, under a GPL-3 license, from the *MDSvis* GitHub repository: github.com/UCLouvain-CBIO/MDSvis. The on-line documentation is available, under a CC-BY-4.0 license, from https://uclouvain-cbio.github.io/MDSvis.

## Acknowledgments

The authors would like to thank Shabnam Zaman and Kim Roelants for providing the *fsap* dataset. This preprint was created using the LaPreprint template (https://github.com/roaldarbol/lapreprint) by Mikkel Roald-Arbøl.

